# ACTIVATION OF CYCLIN-DEPENDENT KINASE 5 BROADENS ACTION POTENTIALS IN HUMAN SENSORY NEURONS

**DOI:** 10.1101/2023.05.31.543017

**Authors:** Manindra Nath Tiwari, Bradford E. Hall, Anita Terse, Niranjana Amin, Man-Kyo Chung, Ashok B. Kulkarni

## Abstract

Chronic pain is one of the most devastating and unpleasant conditions, associated with many pathological conditions. Tissue or nerve injuries induce comprehensive neurobiological plasticity in nociceptive neurons, which leads to chronic pain. Recent studies suggest that cyclin-dependent kinase 5 (CDK5) in primary afferents is a key neuronal kinase that modulates nociception through phosphorylation-dependent manner under pathological conditions. However, the impact of the CDK5 on nociceptor activity especially in human sensory neurons are not known. To determine the CDK5-mediated regulation of human dorsal root ganglia (hDRG) neuronal properties, we have performed the whole-cell patch clamp recordings in neurons dissociated from hDRG. CDK5 activation induced by overexpression of p35 depolarized the resting membrane potential and reduced the rheobase currents as compared to the uninfected neurons. CDK5 activation evidently changed the shape of the action potential (AP) by increasing AP rise time, AP fall time, and AP half width. The application of a prostaglandin E2 (PG) and bradykinin (BK) cocktail in uninfected hDRG neurons induced the depolarization of RMP and the reduction of rheobase currents along with increased AP rise time. However, PG and BK applications failed to induce any further significant changes in addition to the aforementioned changes of the membrane properties and AP parameters in the p35-overexpressing group. We conclude that CDK5 activation through the overexpression of p35 in dissociated hDRG neurons broadens AP in hDRG neurons and that CDK5 may play important roles in the modulation of AP properties in human primary afferents under pathological conditions, contributing to chronic pain.

## 1. INTRODUCTION

Chronic pain affects about 20% of adults in the United States and for its effective treatment, the development of novel efficacious therapies without addiction liability are urgently required ^44^. With ongoing pathological pain signaling, neuroplastic changes can occur that give rise to hyperalgesia and allodynia^19^. Peripheral sensitization, for instance, can arise with transcriptional and post-translational modifications downstream of inflammation. This generally includes the activation of protein kinases such as PKA, PKC, and Cyclin-dependent kinase 5 (Cdk5) in primary afferent neurons^26^. These protein kinases can, in turn, phosphorylate ion channels involved in pain signaling and then can lower their threshold of activation, affect channel desensitization, or enhance channel trafficking to the plasma membrane. Of these protein kinases, Cdk5 is unusual in that it shares a high degree of homology with other Cdk’s that predominately regulates the cell cycle instead of being active in post-mitotic neurons ^34^. In addition, Cdk5 activity is neither induced by a cyclin, as typical with other Cdk family members, nor with a second message molecule as seen with PKA, PKC, and the Ca^2+^/calmodulin-dependent protein kinase CaMKII. Instead, Cdk5 activity is regulated through the expression of its regulatory subunit p35, which binds with Cdk5 through a cyclin box. The upregulation of p35 occurs in the dorsal root ganglia following peripheral inflammation of the rodent hindpaw via carrageenan^26^ or complete Freuend’s adjuvant^42^ or neuropathic injury by spinal nerve ligation^12^. Further studies have shown that inflammatory cytokines including tumor necrosis factor α, transforming growth factor β, and nerve growth factor can trigger ERK1/2 activation to subsequently boost p35 expression and ultimately lead to increased Cdk5 activity^34, 37, 38^.

Cdk5 hyperactivity in primary afferent neurons has consequently been implicated with increased pain hypersensitivity. The transgenic overexpression of p35 alone can, in effect, promote both thermal and mechanical pain in mice^26, 27^. This has been partially attributed to Cdk5 mediated phosphorylation of pain transducing ion channels such as the thermosensor transient receptor potential vanilloid subtype 1 (TRPV1)^25^, the chemosensor transient receptor potential ankyrin subtype 1^12, 14, 32^, and the purinergic receptor P2X2^4^. Phosphorylation by Cdk5 can then essentially potentiate pain signaling by curbing channel desensitization in TPRV1, for example^15^. The dysregulated trafficking of voltage-gated ion channels has also associated with chronic pain, where Cdk5 is known to phosphorylate both Cav3.2, to cause enhanced trafficking to the plasma membrane, and CRMP2, which affects the trafficking of Nav1.7 and Cav2.2^11, 22^.

Pathological pain conditions are accompanied by altered the electrophysiological properties (shape of action potential and intrinsic excitability)^24, 40^. The electrical properties of neurons including action potential, spike width, and spike threshold are tightly regulated by the ionic gradients, ion channels activation/deactivation time and their state of phosphorylation/dephosphorylation^21, 35^. Despite the association of upregulated Cdk5 activity with hyperalgesic conditions, the role of Cdk5 in the modulation of human sensory neuronal properties has not been fully elucidated. Recently, electrophysiological recording of dissociated hDRG neurons has been possible through organ-donor networks^5, 13^. Single nucleus sequencing shows that *CDK5* and its activator p35 (*CDK5R1*) are widely expressed in hDRG neurons^23^. So, an overall assessment of the effects of Cdk5 hyperactivity and other pain associated protein kinases on hDRG neurons has currently become feasible. In this study, to extend our previous preclinical studies on the roles of Cdk5/p35 in pain, we determined the effects of Cdk5 activation on hDRG neuronal firing. We aimed to determine the effects of exogenous overexpression of p35 on the changes in the excitability, action potential properties, and responses to inflammatory mediators in dissociated human DRG neurons.

## 2. MATERIALS AND METHODS

### 2.1. Human subjects

DRGs used in this study were acquired from seven cadaveric donors with the informed consent of the next of kin through Anabios (San Diego, CA). AnaBios Corporation’s procurement network includes only US-based Organ Procurement Organizations and Hospitals. Policies for donor screening and consent are the ones established by the United Network for Organ Sharing (UNOS). Organizations supplying human tissues to AnaBios follow the standards and procedures established by the US Centers for Disease Control (CDC) and are inspected biannually by the DHHS. Distribution of donor medical information complies with HIPAA regulations regarding patient privacy. All transfers of donor organs to ANABIOS are fully traceable and periodically reviewed by US Federal authorities. We obtained approval for carrying out these studies from the National Institutes of Health (NIH) Office of Human Subjects Research Protection (OHSRP) and Biosafety Committee, Bethesda, MD, USA. All methods were performed in accordance with the guidelines and regulations approved by NIH Biosafety Committee, Bethesda, MD, USA.

Table 1 shows the demographics of the donors used for the study. AnaBios generally obtains donor organs/tissues from adults aged 18 to 60 years old. Some donors may be trauma victims, but the donor pool did not include human immunodecifiency virus, donors after cardiac death, hepatitis B virus, ongoing infection, hepatitis C virus, downtime > 20min, methicillin-resitant Staphylococcus aureus, and positive blood cultures.

**Table 1.**
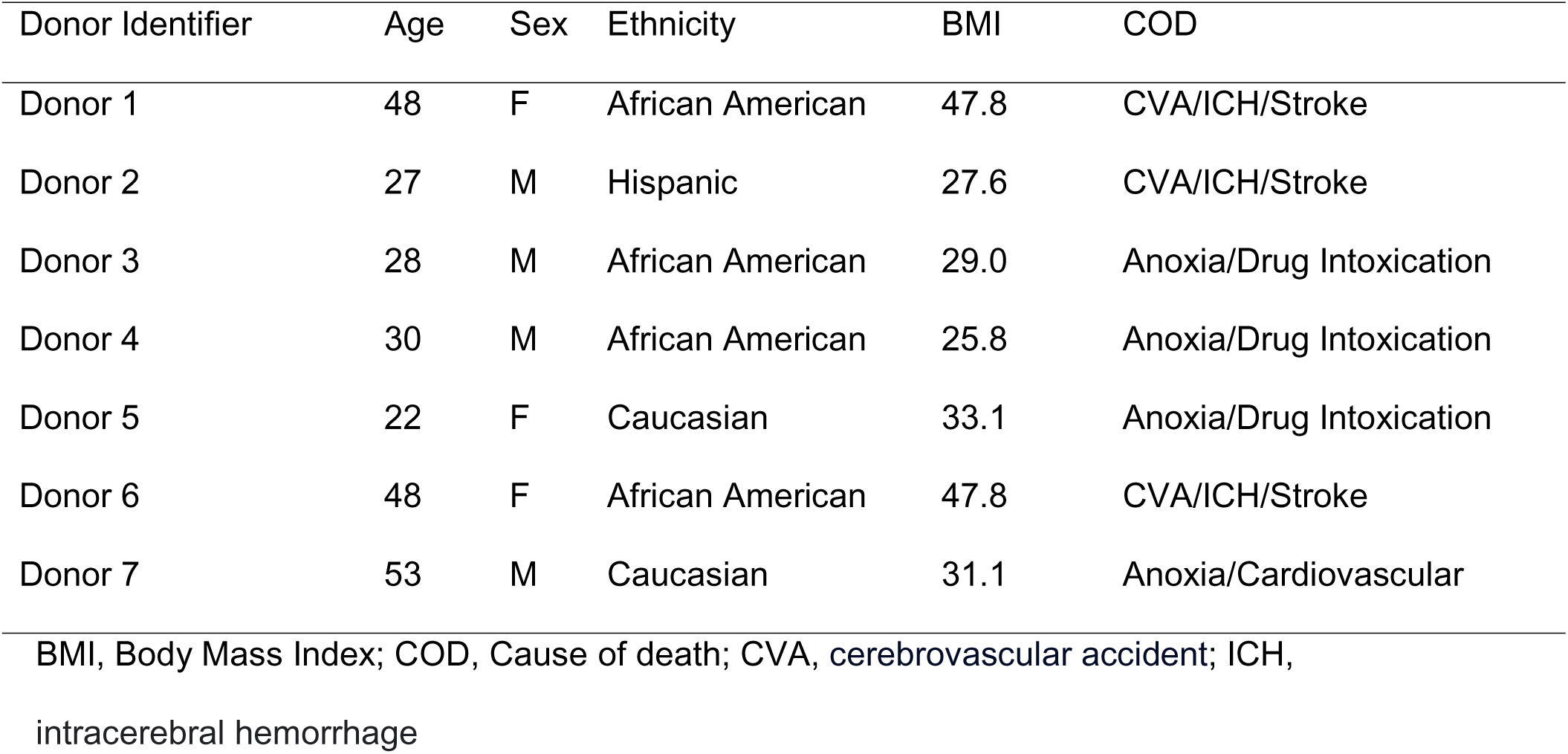
Donor information

| Donor Identifier | Age | Sex | Ethnicity | BMI | COD |
| --- | --- | --- | --- | --- | --- |
| Donor 1 | 48 | F | African American | 47.8 | CVA/ICH/Stroke |
| Donor 2 | 27 | M | Hispanic | 27.6 | CVA/ICH/Stroke |
| Donor 3 | 28 | M | African American | 29.0 | Anoxia/Drug Intoxication |
| Donor 4 | 30 | M | African American | 25.8 | Anoxia/Drug Intoxication |
| Donor 5 | 22 | F | Caucasian | 33.1 | Anoxia/Drug Intoxication |
| Donor 6 | 48 | F | African American | 47.8 | CVA/ICH/Stroke |
| Donor 7 | 53 | M | Caucasian | 31.1 | Anoxia/Cardiovascular |

BMI, Body Mass Index; COD, Cause of death; CVA, cerebrovascular accident; ICH, intracerebral hemorrhage

Donor DRGs (T1-T12, L1-L5, S1) from males and females were harvested using AnaBios’ proprietary surgical techniques and tools shipped to AnaBios via dedicated couriers. Upon arriving at AnaBios, each set of DRGs was assigned a unique identifier number that is reproduced on all relevant medical history files, data entry forms and electronic records. DRG were obtained from 5 donors.

### 2.2. Dissociation of human DRG neurons

The DRGs were then further dissected in cold proprietary neuroplegic solution to remove all connective tissue and fat. The ganglia were dissociated following the methods previously described^5^.

After removing all connective tissue and fat, DRG were enzymatically digested at 37°C for 2 hours using AnaBios’ proprietary mixture of enzymes. After washing, the neurons were resuspended in DMEM/F12 (Lonza; Allendale, NJ) containing 1% horse serum (Thermo Fisher Scientific; Rockford, IL). Ganglia were gently triturated through the fire-polished tip of a sterile glass Pasteur pipette. Dissociated cells were plated on glass coverslips coated by poly-D-lysine. The isolated neurons put in culture in DMEM F-12 (Gemini Bio-Products CAT#: 900-955. Lot# M96R00J) supplemented with Glutamine 2 mM, Horse Serum 10% (Invitrogen #16050-130), hNGF (25 ng/ml) (Cell Signaling Technology #5221LF), GDNF (25 ng/ml) (ProSpec Protein Specialist #CYT-305) and Penicillin/Streptomycin (Thermo Fischer Scientific #15140-122).

### 2.3. Viral infection of hDRG neurons

To confirm expression of the Cdk5 activator p35 before packaging into HSV, an expression vector with an untagged human p35, pCMV-p35 (gift from Dr. Li-Huei Tsai, Addgene plasmid # 1347, Watertown, MA), was transfected into Neuro 2a cells using Neuro-2a Cell Avalanchetransfection reagent (EZ Biosystems, College Park, MD). The expression of p35 was confirmed through Western blot, performed as previously described^33^. After the confirmation of the p35 expression in Neuro 2a cells, pCMV-p35 was cloned into pDONR221 Gateway entry vector (Epoch Life Science; Sugar Land, TX) and packaged into a herpes simplex virus 1 (HSV-1) vector (Dr. Rachael Neve of the Gene Delivery Technology Core, Massachusetts General Hospital). Replication-deficient virus was packaged via the amplicon system and purified on a sucrose gradient. In the HSV vector, an IE4/5 promoter drives p35 expression while the mCMV promoter drives the green fluorescent protein (GFP).

The cultures were infected by either HSV-p35 or HSV-GFP at a multiplicity of infection (MOI) of 3. Uninfected neurons were also assessed to estimate the effects of viral infection. The neurons were used for recordings 1-4 days after infection.

### 2.4. Whole-cell current clamp recordings

Whole-cell patch-clamp recordings were conducted in hDRG neurons 3 to 6 days after dissociation (Fig. 1A). The recordings were performed at room temperature (∼ 23 °C) using HEKA EPC-10 amplifier. Data were acquired on a Windows-based computer using the PatchMaster program (v2x90.4; HEKA Electronics). Pipettes (1.5 -3.0 MΩ) (Warner Instruments #64-0792) were fabricated from 1.5 mm borosilicate capillary glass using a Sutter P-97 puller. The pipette was filled with a solution containing 110 mM K-gluconate, 20 mM KCl, 8 mM NaCl, 4 Mg-ATP, 10 mM EGTA, and 10 mM HEPES (pH 7.3 adjusted with KOH; 280±5 mOsm). Neurons on Corning glass coverslips (Thomas Scientific #354086) were transferred to a RC-26GLP recording chamber (Warner Instruments #64-0236) containing 0.5 ml standard external solution containing 145 mM NaCl, 3 mM KCl, 2 mM CaCl_2_, 1 mM MgCl_2_, 10 mM HEPES, 10 mM dextrose (pH 7.4 adjusted with NaOH; 300±5 mOsm). Extracellular solution exchange was performed with rapid exchange perfusion system (flow rate 0.5-1 ml/min) (Warner Instruments #64-0186). Neurons for recordings were selected based on smoothness of the membrane. Neurons were held at a resting membrane potential (RMP). Signals were filtered at 3 kHz, sampled at 10 kHz. Once whole-cell access was obtained the cell was allowed an equilibration time of at least 5 min. Neurons showing series resistance >15 MΩ were excluded from further recordings. Unless otherwise indicated, all chemicals were purchased from Sigma-Aldrich.

**Figure 1.**
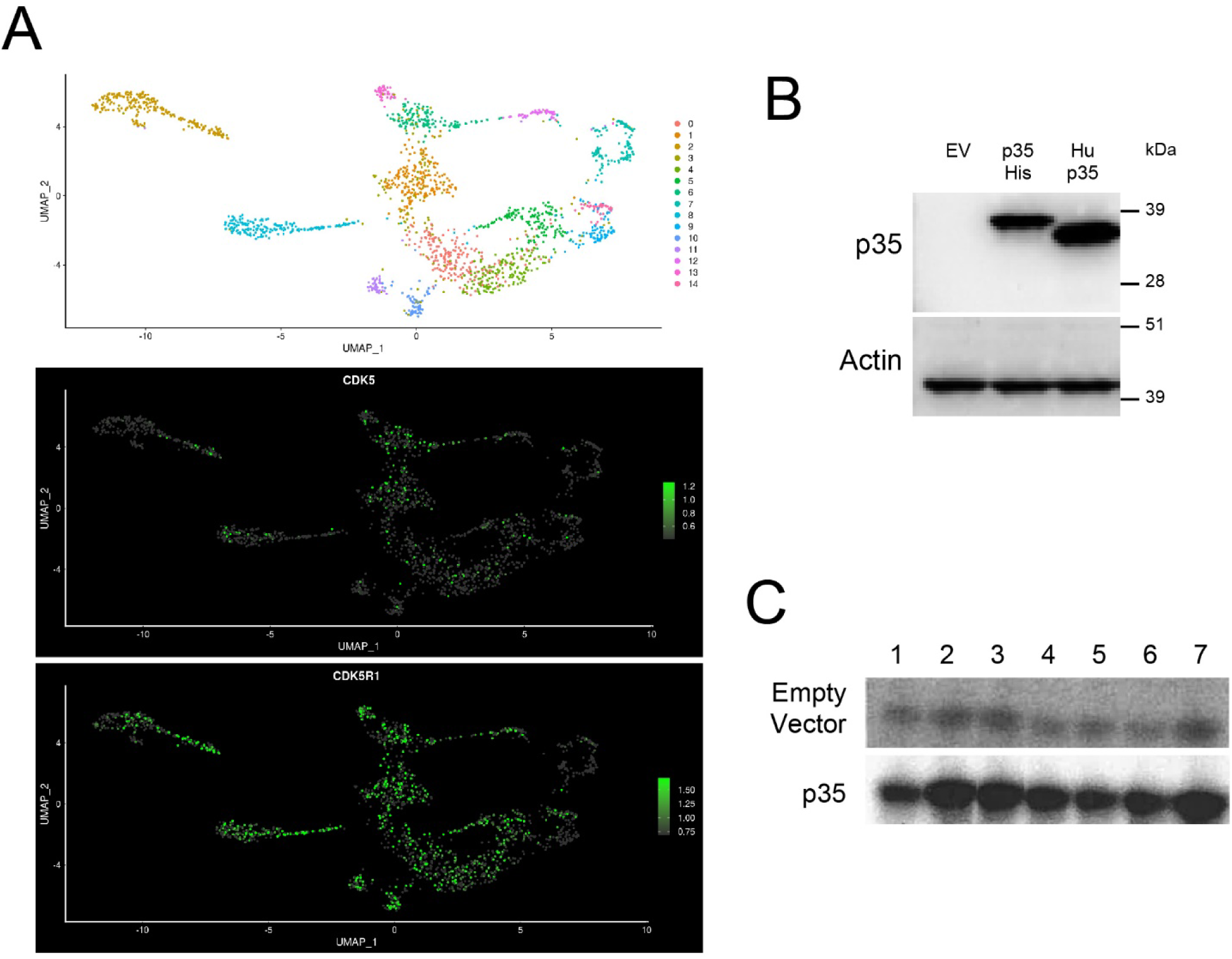
Expression and experimental activation of CDK5 in human sensory neurons. **A.** Single nucleus RNAseq data from hDRG showing the expression of *CDK5* and *CDK5R1* (encoding p35). Data are derived from Nguyen et al., 2021^23^. **B.** A human p35 expression vector was transfected into N2a cells. An empty vector (EV) and a His-tagged p35 vector were used as controls. Western blot for p35 was performed before sending the expression vector to be packaged into HSV. A 35 kDa band corresponding to the untagged human p35 was detected by Western. **C.** SH-SY5Y cells were differentiated and then transfected with a His-tagged p35 expression vector or an empty vector. Next, cells were treated with the following inflammatory mediators: Lane 1, non-treated; lane 2, 100nM bradykinin (BK) + 1µM PGE_2_ (PG); lane 3, 10µM BK + 10µM PG; lane 4, 100nM BK + 1µM PG + 10µM 5HT + 10µM histamine; lane 5, 100nM BK + 1µM PG + 1mM 5HT + 1mM histamine; lane 6, 10µM BK + 10µM PG + 10µM 5HT + 10µM histamine; lane 7, 10µM BK + 10µM PG + 1mM 5HT + 1mM histamine. CDK5/p35 was immunoprecipitated and kinase activity was evaluated by the level of phosphorylated P^32^ histone H1, a substrate of cyclin-dependent kinases.

### 2.5. Measurement of rheobase

Once the cell under recording stabilized, the rheobase of single action potentials was assessed. Step currents in 20 ms duration were delivered every 10 sec. The amplitude of currents was incrementally increased until action potential can be faithfully induced. The step increment was 20 pA if the current amplitude was <1000 pA and 50 pA if the current amplitude was >1000 pA. The minimum current needed for stable action potential induction was documented as rheobase. After measuring rheobase in baseline, 1μM prostaglandin E2 and 100nM bradykinin were applied through bath superfusion (Fig. 1B). The rheobase current was measured after five minutes of exposure.

### 2.6. Measurements of CDK5 activity *in vitro*

To test different inflammatory cocktails for induction of Cdk5 activity, SH-SY5Y cells were differentiated as described in the literature^30^. SH-SY5Y cells were transfected with either an empty vector or with a vector expressing His-tagged human p35 (gift from Dr. Harish Pant) using Lipofectamine 3000 (Life Technologies, Carlsbad, CA). Two days after the transfection, cells were treated with various inflammatory mediators. CDK5 activity was measured as previously described^34^.

### 2.7. Data analysis

In addition to the RMP and rheobase, the kinetics of single action potentials evoked by the rheobase currents were measured (Fig. 1C). Data were expressed as mean±SEM. Kruskal Wallis test (GraphPad Prism v9.0.0) with Dunn’s test multiple comparisons or Wilcoxon signed-rank test were used to determine the statistical significance (α=0.05) as specified in the figure and table legends. The numbers in the results represent the number of neurons regardless of the donors.

## 3. RESULTS

### CDK5 is widely expressed in human sensory neurons and activated by inflammatory mediators

Recent single nucleus RNAseq of human DRG neurons showed that *CDK5* and *CDK5R1*, encoding p35, are expressed in almost every cluster, including peptidergic and non-peptidergic nociceptors^23^ (Figure 1A).

To determine the impact of CDK5 activation on the properties of human sensory neurons, we would take advantage of the fact that p35 overexpression alone produces hyperalgesia in rodents. To achieve p35-induced activation in hDRG neurons, we first generated a p35- expressing HSV vector. The HSV works well for gene delivery into the dissociated DRG neurons^41^. After, we confirmed the protein expression of p35 after transfection by the expression vector (Figure 1B), the coding sequence of an untagged human p35 was then packaged into HSV. This vector was used for dissociated DRG neurons in the following experiments.

Since inflammation of the hindpaw causes increased Cdk5 activity in DRG in rats, we would test if inflammatory mediators increase CDK5 activity in human neuronal cells. We used differentiated SH-SY5Y cells for evaluating CDK5 activity following transfection with either an empty plasmid or with a His-tagged human p35 expression vector. As shown in Figure 1C, 100nM bradykinin (BK) and 1µM prostaglandin E_2_ (PG) showed robust inductions of CDK5 activity, both with and without additional transfection of p35. Therefore, the mixture of these two mediators (PG/BK) was used in the following experiments.

### p35 overexpression tends to depolarize resting membrane potential and to reduce the rheobase currents in hDRG neurons

To determine the changes of the excitability of sensory neurons induced by the activation of CDK5, we infected dissociated hDRG neurons using HSV encoding p35. Neurons infected by HSV encoding GFP served as a control group. To control the effects of HSV transfection, we also tested the uninfected (UI) group. Data were obtained from 10 UI neurons, 9 GFP-expressing neurons, and 9 p35-expressing neurons. By using whole-cell current clamp recordings (Figure 2A), we first assessed resting membrane potential (RMP). Three groups showed significant differences in RMP (p= 0.044 in Kruskal Wallis test). In pair-wise comparisons, p35 overexpression significantly depolarized the resting membrane potential (p= 0.038 in Dunn’s test; Figure 3A) compared to uninfected control groups but not to GFP group (p= 0.86 in Dunn’s test; Figure 3A). Then, we assessed rheobase current by injecting progressively increasing current steps. Three groups showed significant differences in rheobase currents (P=0.033 in Kruskal Wallis test). In pair-wise comparisons, the overexpression of p35 significantly reduced the rheobase current in p35 transfected neurons compared to uninfected control groups. (p= 0.027 in Dunn’s test; Figure 3B). The GFP-infected group also showed a tendency of decreasing rheobase currents, which did not reach statistical significance (p= 0.48 in Dunn’s test; Figure 3B). These results suggest that the activation of CDK5 induces intrinsic excitability by reducing the rheobase current and depolarizing the resting membrane potential of hDRG neurons.

**Figure 2.**
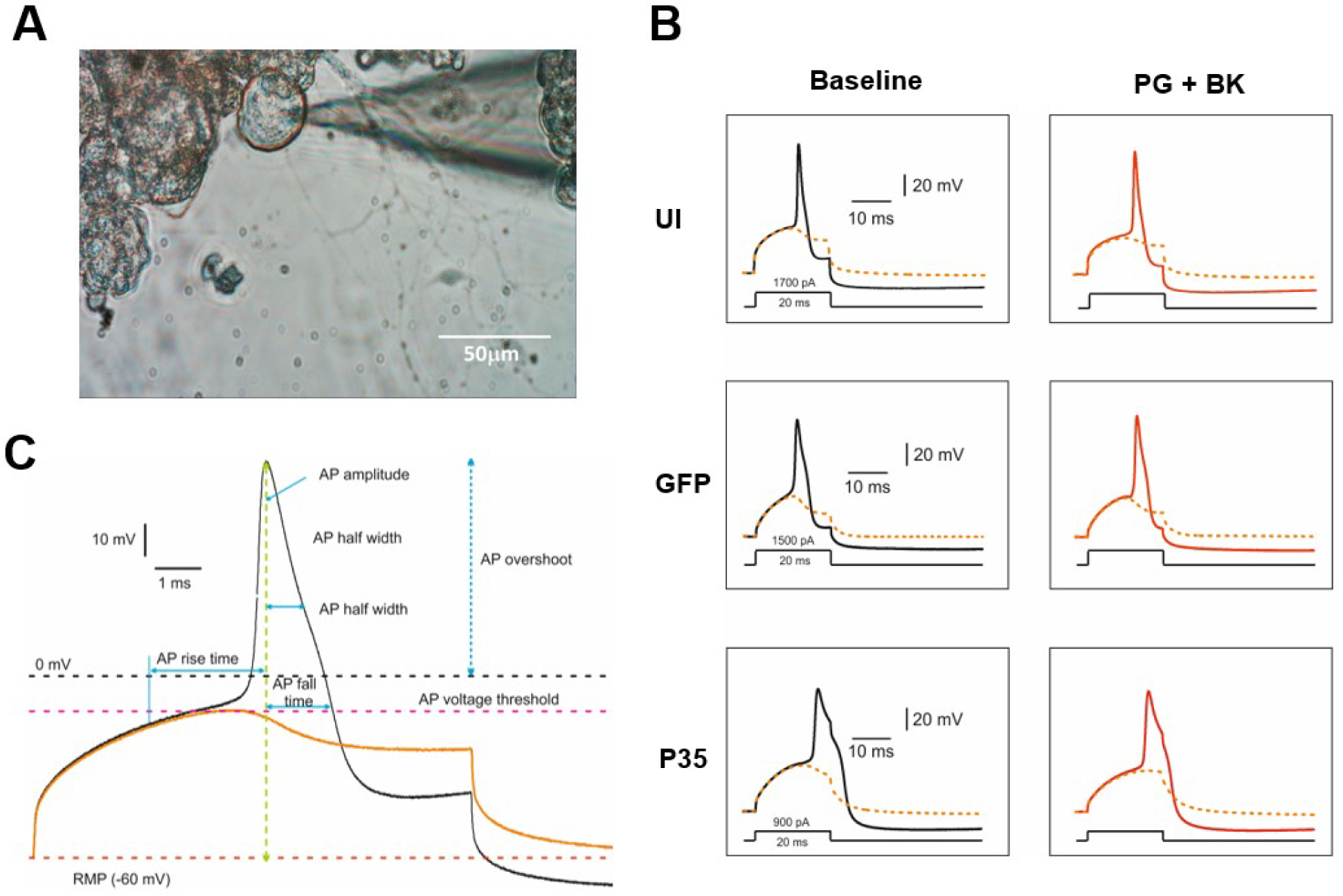
Action potential (AP) measurements in human dorsal root ganglia (hDRG) neurons. **A.** Representative image showing the patched hDRG neurons under the inverted microscope. **B.** A representative AP trace (***black***) evoked at rheobase and a superimposed subthreshold trace without AP (***dashed orange trace***) in three groups (uninfected control; UI), (GFP control; GFP) and (p35 transfected; p35 hDRG neurons) before (baseline) and after perfusion with prostaglandin E2 (1μM) and bradykinin (100nM) (***red trace***). **C.** A representative enlarged AP trace depicting AP measurements. **Rheobase:** Minimal required threshold current to evoke an AP by an incrementing series of depolarizing pulse (incremental step; 50 pA and duration; 20 milliseconds), AP voltage threshold and resting membrane potential **(RMP)** are shown as ***dashed pink line*** and ***dashed red line*** respectively. **AP rise time**: AP rise to the peak, **AP amplitude**: AP peak height from RMP, **AP overshoot:** AP amplitude from zero membrane potential, **AP fall time:** AP peak back to the threshold value, **AHP**: (afterhyperpolarization) amplitude peak amplitude RMP. **AP half width**: AP duration at half the AP amplitude.

**Figure 3.**
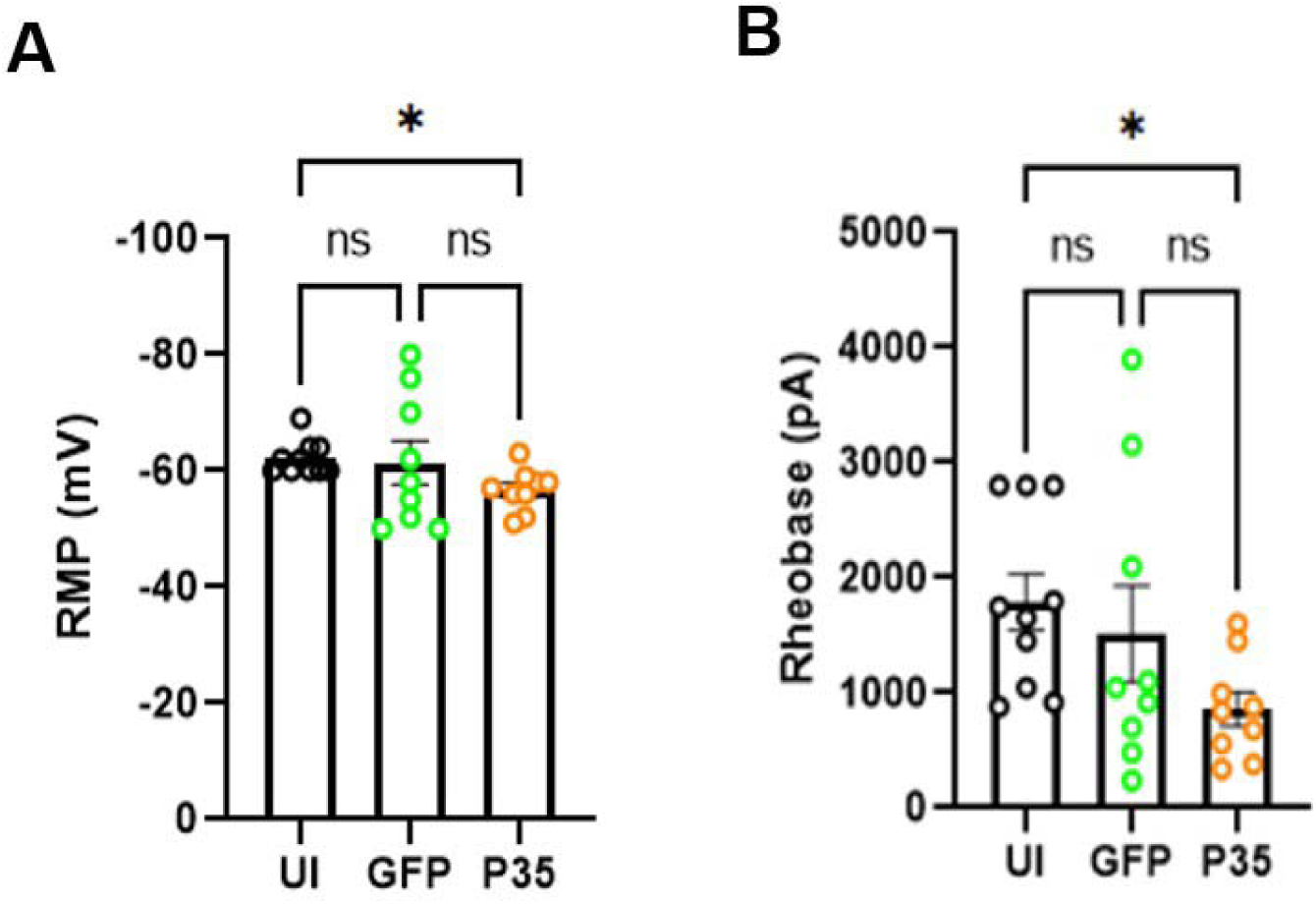
p35 over expression alters the RMP and rheobase current of hDRG neurons. Bar diagrams (mean ± SEM) showing RMP (A), and rheobase current (B) in neurons from UI (*n=10; **black circle***), GFP (*n=9; **green circle***) and p35 (*n=9; **orange circle**)* groups. Statistical comparisons were performed using Kruskal Wallis test followed by Dunn’s multiple comparisons test. \**p* < 0.05, ns (non-significant).

### p35 Overexpression changes the shape of action potential in hDRG neurons

We determined the changes in the AP shape using the parameters described in Figure 2C. Three groups showed significant differences in AP rise time (P=0.003 in Kruskal Wallis test). In posthoc pair-wise comparisons, the overexpression of p35 significantly increases the AP rise time compared to GFP (p= 0.011 in Dunn’s test; Figure 4A, C) and uninfected control groups (p= 0.009 in Dunn’s test; Figure 4A). Three groups showed significant differences in AP fall time (P=0.026 in Kruskal Wallis test). In the multiple comparisons, the overexpression of p35 significantly increases the AP fall time compared to GFP group (p= 0.03 in Dunn’s posthoc test ; Figure 4B). Consistently, three groups showed significantly different AP half width (p= 0.003 in Kruskal Wallis test), and the posthoc test showed that the overexpression of p35 significantly increases the AP half width compared to GFP (p= 0.011 in Dunn’s test; Fig. 4C) and uninfected control groups (p= 0.007 in Dunn’s test; Figure 3C).

**Figure 4.**
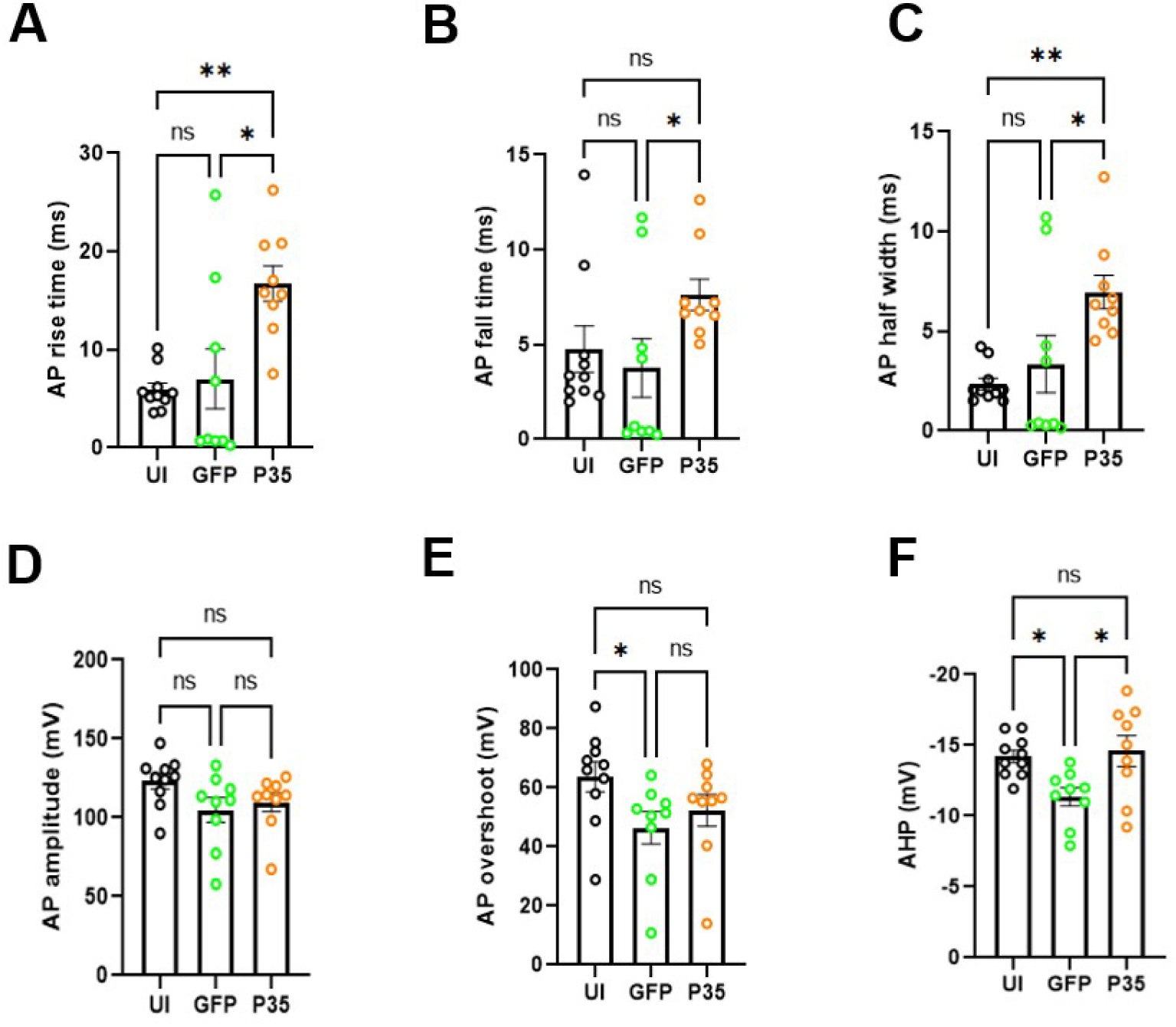
p35 over expression alters the AP properties in hDRG neurons. Bar diagrams (mean ± SEM) showing AP rise time (A), AP fall time (B), AP half width (C), AP amplitude (D), AP overshoot (E), and AHP (F) in neurons from UI (*n=10; **black circle***), GFP (*n=9; **green circle***) and p35 (*n=9; **orange circle**)* groups. Statistical comparisons were performed using Kruskal Wallis test followed by Dunn’s multiple comparisons test. \**p* < 0.05, \*\**p* < 0.01, ns (non-significant).

Three groups did not show differences in AP amplitude (P= 0.067 in Kruskal Wallis test; Figure 4D). Although AP overshot was significantly different among the three groups (P= 0.024 in Kruskal Wallis test) posthoc pair-wise comparisons showed a significant difference only between UI and GFP (p= 0.02 in Dunn’s test; Figure 4E). The afterhyperpolarization potential (AHP) was significantly different among the three groups (P= 0.01 in Kruskal Wallis test) and the posthoc test showed that AHP in the GFP group was significantly reduced compared to the UI (p= 0.04 in Dunn’s test; Figure 4F) and the p35 groups (p= 0.02 in Dunn’s test; Figure 3F). The changes in the shape of action potential suggest that activation of CDK5 by p35 overexpression broadens the action potential in hDRG neurons.

### p35 overexpression does not enhance the responsiveness of hDRG neurons to prostaglandin E2 and bradykinin

In the second part of our study, we investigated the effect of inflammatory mediators (PG/BK) on the electrophysiological properties of hDRG neurons in p35-transfected, GFP-transfected, and uninfected groups.

After assessing RMP, rheobase currents, and AP properties under baseline condition, PG/BK was applied to the bath for five minutes and AP was evaluated again. PG/BK significantly depolarized the RMP in UI (p= 0.03 in Wilcoxon signed rank test Figure 5A, upper panel). However, PG/BK exposure showed a tendency of depolarization in GFP, but was not significantly different (p= 0.14; in Wilcoxon signed rank test Figure 5A, upper panel). p35 group did not show a difference either after PG/BK application (p= 0.41; in Wilcoxon signed rank test Figure 5A, upper panel). The percent change of RMP was significantly different among the three groups (P= 0.034 in Kruskal Wallis test figure 5A, lower panel) but posthoc pair-wise comparisons showed no significant changes (figure 5A, lower panel). Next, we assessed the rheobase current before and after PG/BK exposure. PG/BK application significantly reduced the rheobase currents compared to baseline in all three groups (UI: p= 0.002; GFP: p= 0.04 and p35: p= 0.02 in Wilcoxon signed rank test; Figure 5B, upper panel). Comparisons of percent changes of rheobase currents among three groups showed a significant difference (P= 0.023 in Kruskal Wallis test Figure 5B, lower panel) and post-hoc pair-wise comparisons showed that p35 group showed a significantly lower change in rheobase currents in response to PG/BK compared to GFP group (p= 0.02 in Dunn’s test; Figure 5B, lower panel).

**Figure 5.**
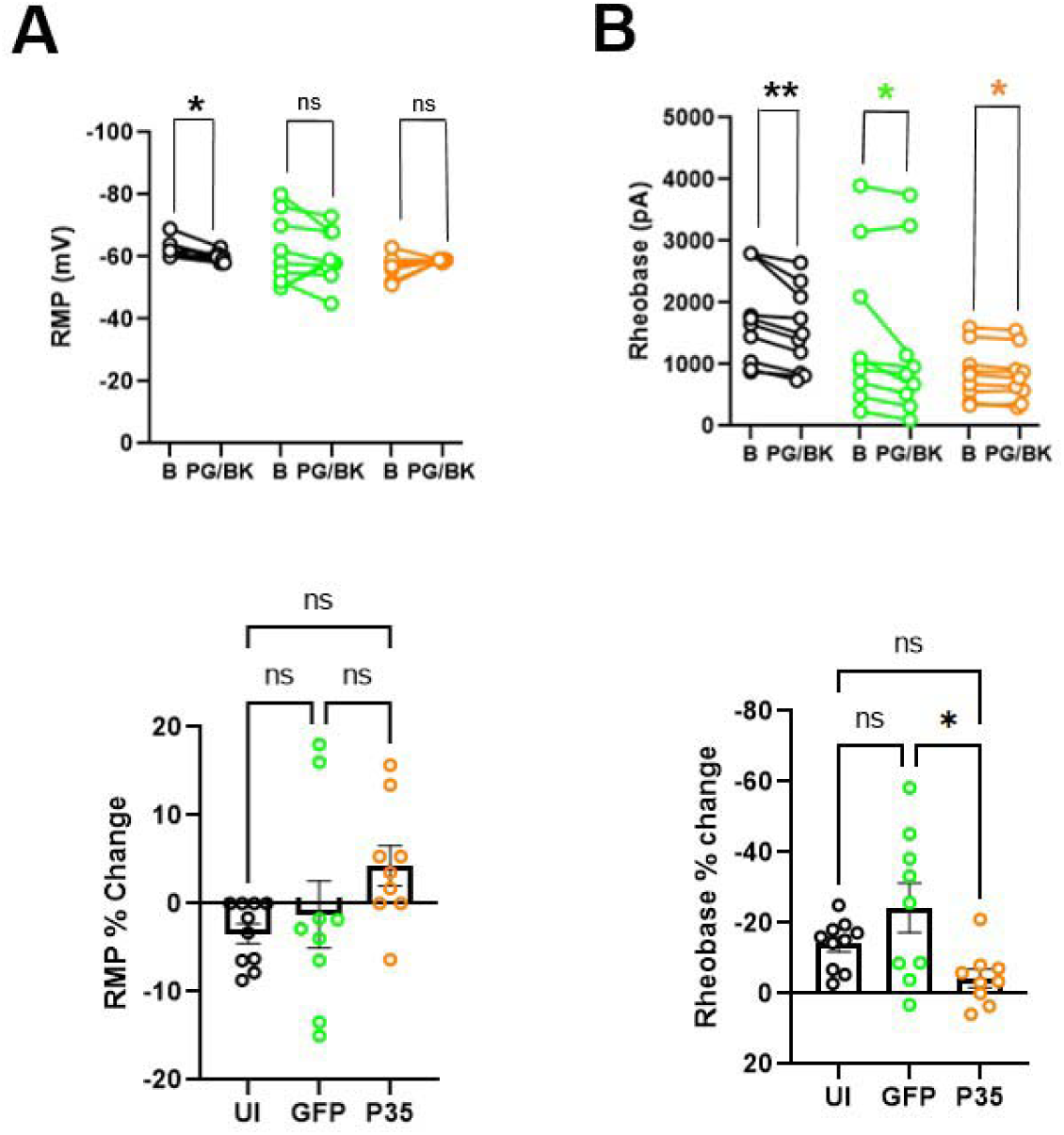
The effect of prostaglandin E2 (PG) and bradykinin (BK) on the RMP and rheobase current of hDRG neurons. Paired dot plot graph depicting (A) RMP, and (B) rheobase current before (baseline; B) and after 1μM prostaglandin E2 and 100 nM bradykinin (PG/BK) in UI (*n=10; **black circle***), GFP (*n=9; **green circle***) and p35 (*n=9; **orange circle**)* neurons. In each section, the upper panel represents absolute value and lower panel represents % change. Statistical comparisons were performed using Wilcoxon signed rank test for paired data analysis and Kruskal Wallis test for percent changes. Dunn’s multiple comparisons test was performed for pairwise comparisons of percent changes. \**p* < 0.05, \*\**p* < 0.01, ns (non-significant).

PG/BK application significantly increases the AP rise time in UI (p= 0.014), but not in GFP (p= 0.3) and p35 group (p= 0.16) (Wilcoxon signed rank test; Figure 6A, upper panel). The percentage changes of the three groups were not significantly different. Other AP parameters were not significantly affected by the PG/BK exposure in any groups (Figure 6B-F).

**Figure 6.**
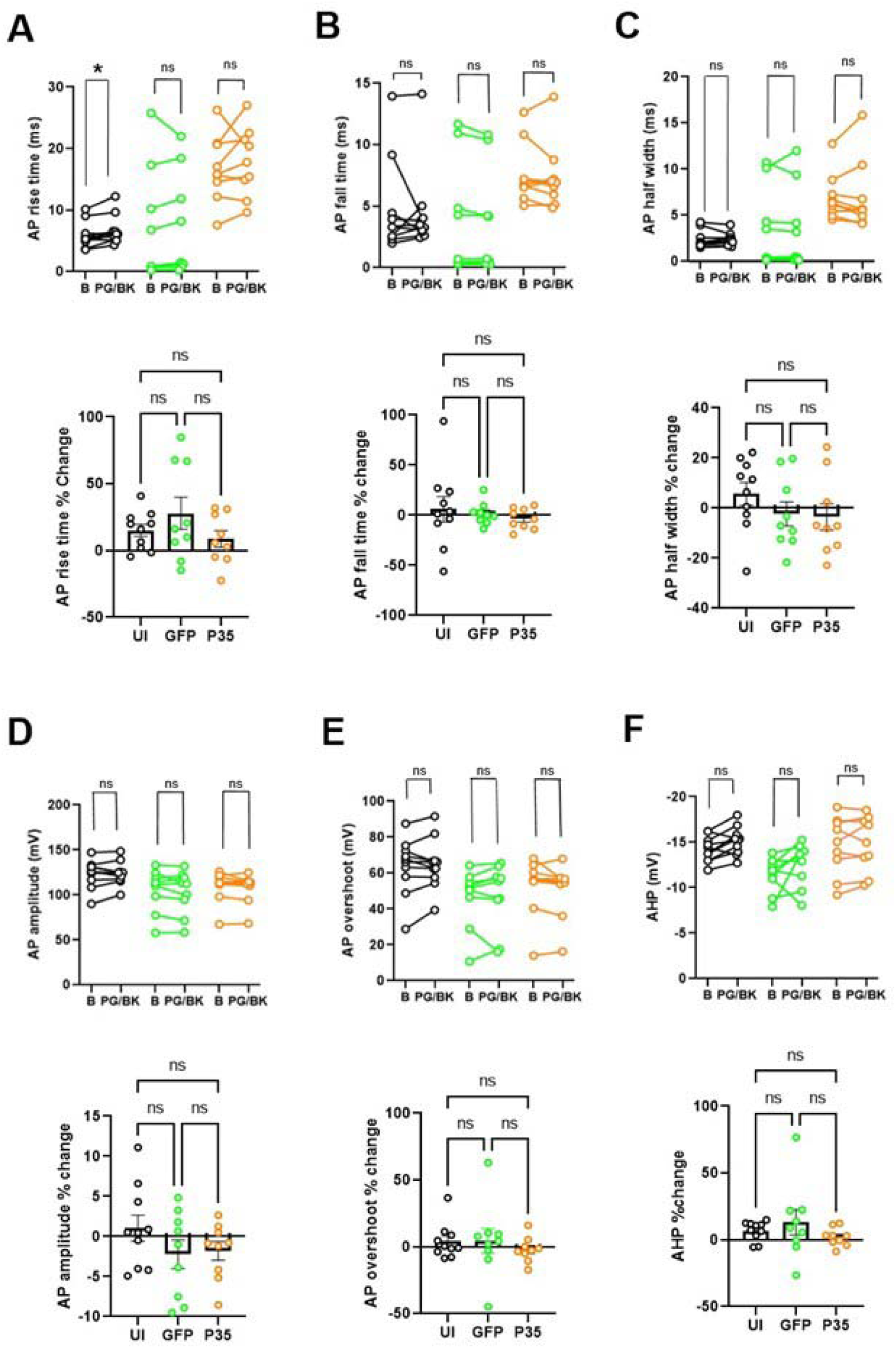
The effect of prostaglandin E2 (PG) and bradykinin (BK) on the AP properties of hDRG neurons. Paired dot plot graph depicting (A) AP rise time, (B) AP fall time, (C) AP half width, (D) AP amplitude, (E) AP overshot, (F) AHP before (baseline; B) and after 1μM prostaglandin E2 and 100 nM bradykinin (PG/BK) in UI (*n=10; **black circle***), GFP (*n=9; **green circle***) and p35 (*n=9; **orange circle**)* neurons. In each section, the upper panel represents absolute value and lower panel represents % change. Statistical comparisons were performed using Wilcoxon signed rank test for paired data analysis and Kruskal Wallis test for percent changes. Dunn’s multiple comparisons test was performed for pairwise comparisons of percent changes. \**p* < 0.05, \*\**p* < 0.01, ns (non-significant).

## 4. DISCUSSION

The contribution of CDK5 in primary afferents has been well established through preclinical models. However, there is no evidence supporting the roles of CDK5 in nociceptor functions in humans. In this study, we have used an electrophysiological approach to investigate the role of CDK5 on the membrane properties and action potential properties of hDRG neuronal culture. To the best of our knowledge, this is the first report demonstrating that the activation of CDK5 influences the AP properties in human sensory neurons. In this study, we designed to test two pathologically relevant conditions to determine the impact of CDK5 activation on the properties of hDRG neurons. First, we mimicked the upregulation of CDK5 activity by the increased expression of p35. In preclinical models, overexpression of p35 alone increases pain sensitivity in mice without producing inflammation or nerve injury^26, 27^, suggesting that p35 overexpression is sufficient to produce hyperalgesia. We achieved this condition by using the HSV vector for overexpressing p35 in dissociated neurons. Second, we mimicked the inflammatory condition under which nociceptors are exposed to inflammatory mediators. Based on our data supporting the activation of CDK5 by PG/BK, we exposed hDRG neurons to PG/BK to assess their effects on electrophysiological properties. Although the large variations of data did not produce significant differences, p35 overexpression showed a tendency to enhance the excitability of hDRG neurons by changing the rheobase current and RMP. Importantly, p35 overexpression altered the AP rise time, AP fall time and AP half-width leading to AP broadening. We acknowledge that large variations of the data limit our ability to be conclusive in membrane properties (RMP and rheobase) and that HSV infection itself apparently affects the properties of hDRG neurons. Nonetheless, the impact of p35 overexpression on the broadening of AP was evident, which strongly supports the idea that CDK5 hyperactivity influences the electrophysiological properties of hDRG neurons.

Acute application of PG/BK in bath altered the membrane properties (rheobase and RMP) and AP properties (AP rise time) in uninfected hDRG neurons. This is consistent with the fact that inflammation enhances Cdk5 activity in DRG of rodents^43^. However, PG/BK application did not produce any significant changes either in membrane properties or AP properties in GFP- and p35-expressing groups. Given the tendency of depolarized membrane potential and decreased rheobase following p35 transfection, we postulate that additional activation of CDK5 by PG/BK following p35 overexpression may be modest and does not produce any further changes. Again, however, the interpretation of the data is limited by the large variation of the data, the impact of HSV infection, and the insufficient sample size.

AP time course represents the ensemble of depolarizing and repolarizing ionic conductances. Intricate balances of voltage-gated sodium, calcium, and potassium channels produce neuronal AP firing. Different inward currents from TTX-sensitive and TTX-resistant Na^+^ channels and Ca^2+^ currents contribute to different phases of AP in DRG neurons^2^. Action potential of nociceptive neurons shows wide action potential with a characteristic hump on the falling phase^7, 9, 20, 28, 39^. Such characteristic is attributable to TTX-resistant Na^+^ channels and voltage-gated Ca^2+^ channels. Large conductance Ca^2+^-activated K^+^ channels also contribute to the duration of AP in non-peptidergic nociceptive DRG neurons^45^. It is not clear what is the functional implication of the wide AP in nociceptors. We speculate that nociceptors may be optimal for the transduction and conduction of low-frequency stimuli. AP broadening should increase Ca^2+^ influx in presynaptic terminals, which can induce more efficient nociceptive synaptic transmissions. The peripheral and central nociceptive system may be optimized for the low-frequency encoding in the transduction and conduction of nociceptive signal to differentiate it from the high-frequency non-nociceptive conduction systems, such as tactile sensation or proprioception.

Importantly, AP width is not static, but the broadening of AP in sensory neurons occurs in multiple pathological conditions through various mechanisms. For example, monocyte chemoattractant protein-1 broadens action potential in DRG neurons through the increase of non-selective cationic currents and the inhibition of voltage-dependent K^+^ currents^33^. Chronic constriction injury of the infraorbital nerve or chemotherapy-induced neuropathy by oxaliplatin leads to an increase in AP widths of trigeminal ganglia neurons, which is likely through the downregulation of Kv4.3^16, 42^. Peripheral axotomy broadens AP mediated by the downregulation of Kv2.1 and Kv2.2^36^. Mutation of Nav1.7 associated with erythromelalgia increases AP width^18^. Our study shows that the activation of CDK5 through p35 overexpression broadens AP in human DRG neurons. Since Cdk5 is upregulated in the DRG after spinal nerve ligation in rats^11^, AP broadening can be associated with the pathological condition likely contributing to neuropathic pain in humans. Therefore our data strongly suggest that CDK5 can play a modulatory role in human primary afferents contributing to chronic pain.

Since we did not evaluate CDK5-induced changes in ionic currents by voltage clamp, we do not have evidence underlying the mechanisms of AP broadening. AP width is predominately determined by Ca^2+^ transients, potassium, and sodium channel activities^1, 6, 17^. The inhibition of Cdk5 activity reduces the TRPV1-mediated Ca^2+^ influx in rat DRG neurons ^25^. Nav1.7 channel activity is regulated by CDK5-dependent phosphorylation of CRMP2 (a binding partner of NaV1.7) in neuropathic pain condition^8^. Cdk5 also phosphorylates and regulates both high voltage-activated N-type Cav2.2^31^ and low voltage-activated T-type Cav3.2^10^ Ca^2+^ channels. CDK5 regulates the voltage-gated Kv2.1 channels through direct phosphorylation, which is crucial to activity-dependent plasticity in intrinsic neuronal excitability^3^. CDK5 also regulates the activity of Kv7.2 channels through its phosphorylation^29^. Our substrate analysis suggested that CDK5 may phosphorylate other voltage-gated ion channels, such as Nav1.9, Cav3.3, Kv6.2, and K_Ca_2.1^14^. Therefore, CDK5 activation may broaden AP in hDRG through the ensemble regulation of the voltage-gated ion channels, which suggests that CDK5 is an important upstream regulator of AP properties in pathological conditions.

In conclusion, CDK5 activation through the exogenous overexpression of p35 in dissociated hDRG neurons broadens AP. Our data suggest that CDK5 plays an important role in the modulation of AP properties in human primary afferents under pathological conditions and contributes to chronic pain.

## 5. Acknowledgements

This work was supported by the National Institutes of Health Grants ZIA-DE-000664-24 to ABK and R01 DE027731 and R35 DE030045 to MKC. We would like to thank Dr. Rachael Neve for preparation of the HSV vectors, Dr. Michaela Prochazkova for help with analyzing inflammatory cytokines, and Drs. Minh Nguyen and Nick Ryba for help analyzing single-nucleus transcriptomic data. We would also like to thank AnaBios, LLC for harvesting DRGs from organ donors, culturing DRG neurons, and performing whole-cell patch-clamp recordings.

